# Variable and reversible opacification process on IOLs for cataract simulation

**DOI:** 10.1101/2023.09.27.559841

**Authors:** Deniz Akyazi, Parviz Zolfaghari, Afsun Sahin, Hakan Urey

## Abstract

Understanding vision through mild and dense cataracts is important for vision scientists and IOL developers. There have been virtual simulators using VR headsets for subjective evaluation of cataractous vision. However, a physical intraocular lens with variable cataracts for objective evaluation is not available. In this study, we developed a reversible process that can be selectively applied to the whole or parts of an IOL to affect the opacification level. We used our eye model and developed a cataractous lens simulator for progressive levels of opacification, which is achieved by applying a reversible chemical procedure on the intraocular lens surfaces. After the lens is fully or partially immersed in acetone, subsequent testing of the lens in distilled water results in a progressive change in opacification level within minutes. We measured the quality of vision by obtaining modulation transfer function curves, transmission, and spectroscopic measurements at different opacification levels. By simulating variable opacification across the IOLs, we tested how vision changes from less dense to more dense cataractous regions in a holographic display system with programmable small exit pupils. All results were consistent with the expected vision degradation caused by natural opacification.

## Introduction

As the first report assessing vision and visual impairments worldwide, the World Report on Vision stated in 2019 that among the 2.2 billion people affected by visual impairment globally, 33% suffer from cataracts [1]. The clouding of the crystalline lens and the presence of visual symptoms, such as decreasing contrast sensitivity and visual acuity, blurred vision, color shift, and photic phenomena, constitute the characteristics of the disease known as cataracts [2]. In the realm of vision research, advancements in optical techniques have contributed significantly to our understanding of various ocular conditions. These innovations, including optical methods for studying and assessing cataracts, glaucoma, and other eye disorders, play a crucial role in comprehending the broader landscape of vision health [3-5].

With today’s technology, cataracts can be treated with a simple surgery; however, some severe complications persist afterward, such as posterior capsule opacification (PCO) or decentration. PCO is one of the most common complications requiring treatment post-operatively in up to 50% of patients [6]. Treatment of PCO can be done with YAG laser capsulotomy.

Although cataracts and PCO are easy to treat and there have been standardized methods to assess the symptoms caused by them, their progression is slow, making them hard to keep track of. People with such complications may not notice them until cataracts or PCOs become severe, significantly decreasing their quality of life.

There have been previous studies that aimed to simulate how cataracts affect vision. Augmented and virtual reality were the common choices in these simulations since these technologies were often used in ophthalmologic studies [2, 7-10]. Unlike the discussed simulators, another study designed an optical device using protein denaturation in egg albumen [11]. This study could successfully mimic the progression of cataracts and their effect on vision, but this progress was irreversible. Once the progress was complete, the device had to be set up from scratch for another cataract mimicking, making this method impractical for further analysis.

In this study, we tried to demonstrate progressive opacification with a simple chemical and reversible process. This process uses basic materials such as acetone and distilled water. We conducted simulations to assess visual outcomes in individuals with cataracts, employing an artificial eye model [12] and a chemical process on intraocular lenses. By repeating the same method on one side of the IOL, it was also possible to assess the vision with a progressive PCO. It would also be possible to demonstrate different cataract types by controlling the degree of opaqueness in specific areas. The influence of varying degrees of cataracts on visual quality was objectively evaluated by acquiring modulation transfer function curves and spectroscopy results. Furthermore, by simulating opacification in a structured fashion, we examined the variations in results within the patient’s pupil, transitioning from regions with lower cataract density to those with higher density, using our holographic display as a testing platform.

## Methods

### Progressive Opacification

We developed a procedure utilizing the reaction between distilled water and acetone to create a reversible and progressive opacification on the IOL (Tecnis monofocal) surface. As discussed, this was done to simulate PCO and cataracts in the human eye and how they affect vision. The procedure consists of the following steps:

1. The IOL is cleaned with isopropanol to remove any previous opacification or dirt.

2. The cleaned IOL is then soaked in distilled water.

3. The wet IOL is directly dipped into pure grade (99.5%) acetone and kept for about 15 seconds.

4. Then, the IOL is put aside to dry for 30 seconds.

5. Finally, once the processed IOL is put into a wet cell, it gradually becomes opaque over time as it stays in the water.

After the IOLis is put into the water, it experiences a continuous buildup of opacity on its surface. This phenomenon persists until the IOL is removed from the medium. Notably, the reversal of this opacification is a straightforward procedure. It can be achieved either by immersing the IOL in acetone, which leads to an immediate and complete reversal, or by soaking it in alcohol, which rapidly eliminates the opacification within seconds.

### Objective Analysis on Vision with Progressive Opacification

The vision through the IOL with various opacification levels was captured using the artificial eye model we designed according to ISO 11979-2 standards [13] using a scleral rigid gas permeable contact lens and an IOL immersed in distilled water [12]. To objectively quantify the degradation of vision, several assessments were done. MTF charts for different opacification levels were found by capturing and analyzing a slanted edge. The transmission throughout a spectrum between 200 nm and 1000 nm was measured using a spectroscope. With a photodiode, the transmission as well as scattering for a light source of 530 nm wavelength were measured. Finally, the color shift was found by analyzing the color histogram of captures through the opacification process.

### MTF Charts

In the first characterization, to understand how the resolution limit is affected by opacification, a slanted edge was captured through the artificial eye model implanted with an opaque cataract and the modulation transfer function analysis was conducted. We converted the corresponding spatial frequency unit of cycles per pixel to cycles per degree to understand the graphs clearly by using the parameters given in Table 1.

**Table 1.** Parameters of the slanted edge capture setup.

|  |  |
| --- | --- |
| **Camera pixel pitch** | 3.45 $\mu\text{m}$ |
| **Distance between chart and IOL** | 100.0 mm |
| **Distance between IOL and camera** | 8.7 mm |

### Transmission and Scattering

One of the biggest known aspects of cataracts, or PCO, is the scattering surface. To first verify and understand the scattering in our cataract mimicking, we utilized a photodiode and measured the transmission through the IOL with the opacification process as a function of time. The setup for this application is given in S1 Fig. In the second step, we put an aperture in front of the photodiode and placed it such that it was at the focal plane of the IOL. This way, we eliminate the scattered light and measure how much of the transmitted light still stays on the optical path. For the light source, we used an LED light source with a 530 nm peak wavelength and collimated it with an ideal lens to create a beam of 2 mm diameter. The light source’s input power was 0.7 mW.

### Spectroscopy

In the next characterization, we utilized spectroscopy and a light source with a spectrum from 200 nm to 1000 nm to see how opacification affects different wavelengths with a setup given in S2 Fig. We added a neutral-density filter in front of the light source to get rid of saturation. After the light passed through the filter, we put the IOL just in front of the spectrophotometer and conducted the measurements. Once again, with an interval of 60 seconds, spectroscopy measurements were taken on the IOL with the proceeding opacification up to 300 seconds.

### Holography and Patterned Opacification

In addition to the characterization, an experiment was carried out to examine the behavior of light transmission and scattering through the opacified intraocular lens in holographic displays. In this step, both the unaltered (original) and opaque states of the IOL were positioned inside the holographic setup. The device was made up of a phase spatial light modulator (SLM), a green LED of 530 nm acting as the light source, and a 4f optical system made up of two lenses that worked as a spatial filter to eliminate unwanted diffraction patterns. A magnifying lens was also incorporated into the system. Holographic images of a Snellen chart were transmitted, through the IOL, and captured by a scientific camera. This was accomplished by submerging an IOL inside a wet cell and placing the scientific camera behind it. The setup used and its details are given in S3 Fig. There were two main steps in the experimental process. Initial holographic images of a Snellen chart were captured while the holographic configuration was set up for the unaltered IOL. A sequence of sequential holographic images was then recorded after the opacification procedure was applied and the IOL was kept soaked in water. In the second step, we also created a patterned opacification on the IOL by clearing some opaque parts with acetone. With this, a circular region about 2 mm in diameter was turned back to transparent, allowing the holographic image to pass through that region. In conclusion, the investigation focused on the light behavior of opaque IOLs and used a holographic setup to simulate the optics of the human eye. The procedure involved taking multiple holographic images of a Snellen chart, first with the IOL as-is and then after performing the opacification on the IOL.

The patterned opacification was tested using holography due to its advantage of pinhole display and based on a previous study done by our group [14]. Since the visual acuity was assessed by them utilizing the pinhole display and sending the holographic image through the non-cataract region of the patient’s eyes, we wanted to understand how well this method performed by repeating it on our artificially opaque IOL surface.

## Results

### General Procedure

The opacification progression through time was first captured for direct observation (Fig 1). In intervals of around 40 seconds, the IOL was taken out of the wet cell and captured on a meter. One can observe that as time passes, the IOL becomes visually opaque, barely transmitting light, and the number behind it gets less visible, eventually not visible at all. The limit of 300 seconds was chosen because, after 300 seconds, the degradation of vision was barely noticeable to the human eye. An example for patterned opacification is given in S4 Fig.

**Fig 1.**
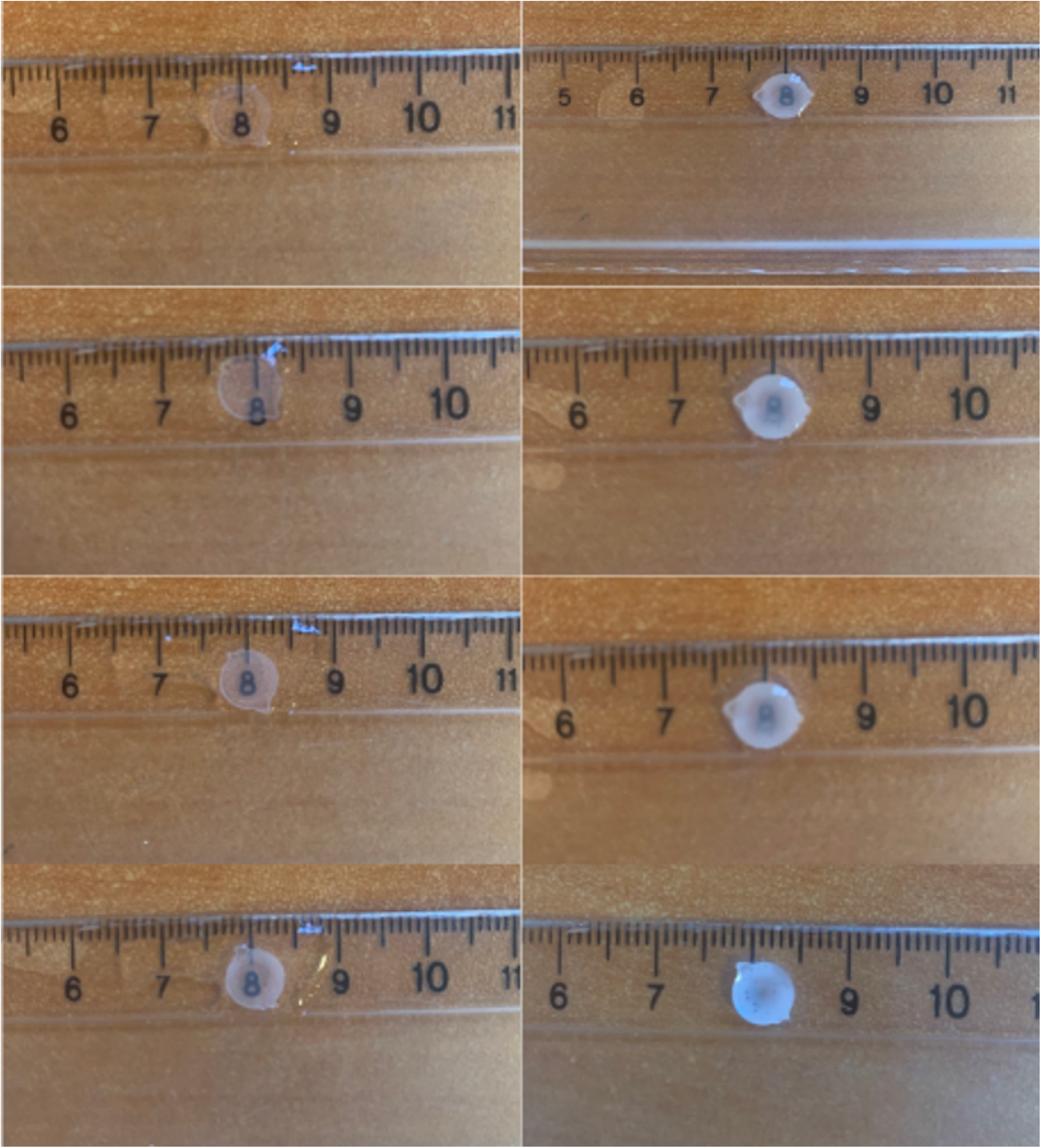
Opacification progress on IOL. Captures of progressive opacification on the IOL after (A) 0, (B) 40, (C) 80, (D) 120, (E) 160, (F) 200, (G) 250, and (H) 300 seconds.

### Vision through the cataracts

Several scenes were captured through the IOL with opaque surfaces. To simulate different scenarios, an office inside, a nighttime courtyard of our university, and a reading chart were captured with different opacification levels.

The first capture discussed is of a scene at the court of our university in the daytime (Fig 2). One of the most important aspects of this capture is that we can clearly observe that the transmitted light decreases as the opacification level increases, turning the scene darker. Apart from the imperfections due to the original IOL, blur starts to occur and increases with opacification. When we pay attention to the lights in the ceiling, even though they are saturated in the first capture with no opacification, as opacification increases, they become saturated and turn yellowish. These results are all known degradations of vision caused by cataracts.

**Fig 2.**
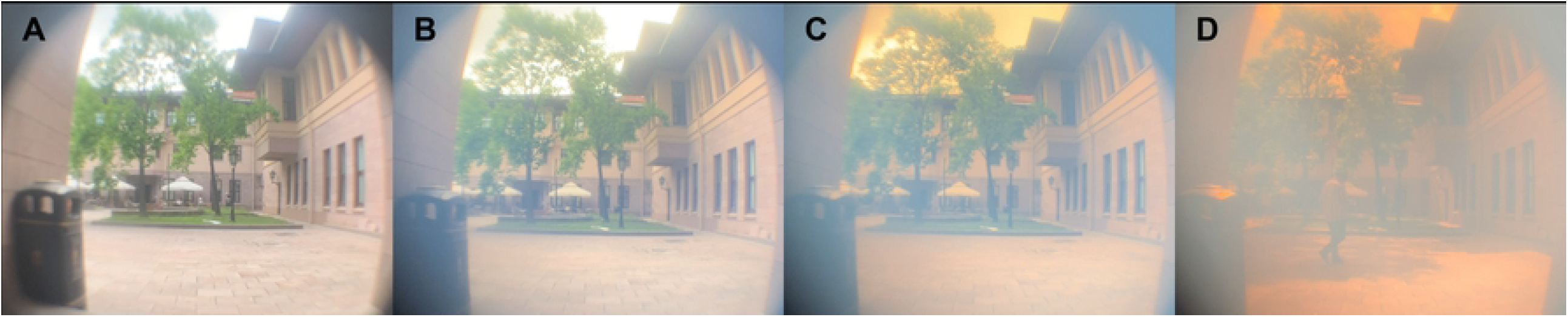
Daytime capture through progressive opacification on the IOL. A court area of a university building is captured through an artificial eye model implanted with different opacification levels on the IOL at (A) 0 seconds, (B) 100 seconds, (C) 200 seconds, and (D) 300 seconds.

The second capture is the night-time capture of a court in our university through different levels of opacification (Fig 3). Similar results can be observed with the captures of the office scene. The light sources are getting unsaturated, and the scene is becoming blurry and darker with a color shift towards red. Another prominent detail is the scattering of the light. When the edges of the capture are observed, we can see that the light coming from the light scene scatters and overflows to the sides, creating a glare like phenomenon, which points out that our opacification not only decreases the transmission of the light but also scatters it. This is another known feature of cataracts, so it further agrees with our simulations.

**Fig 3.**
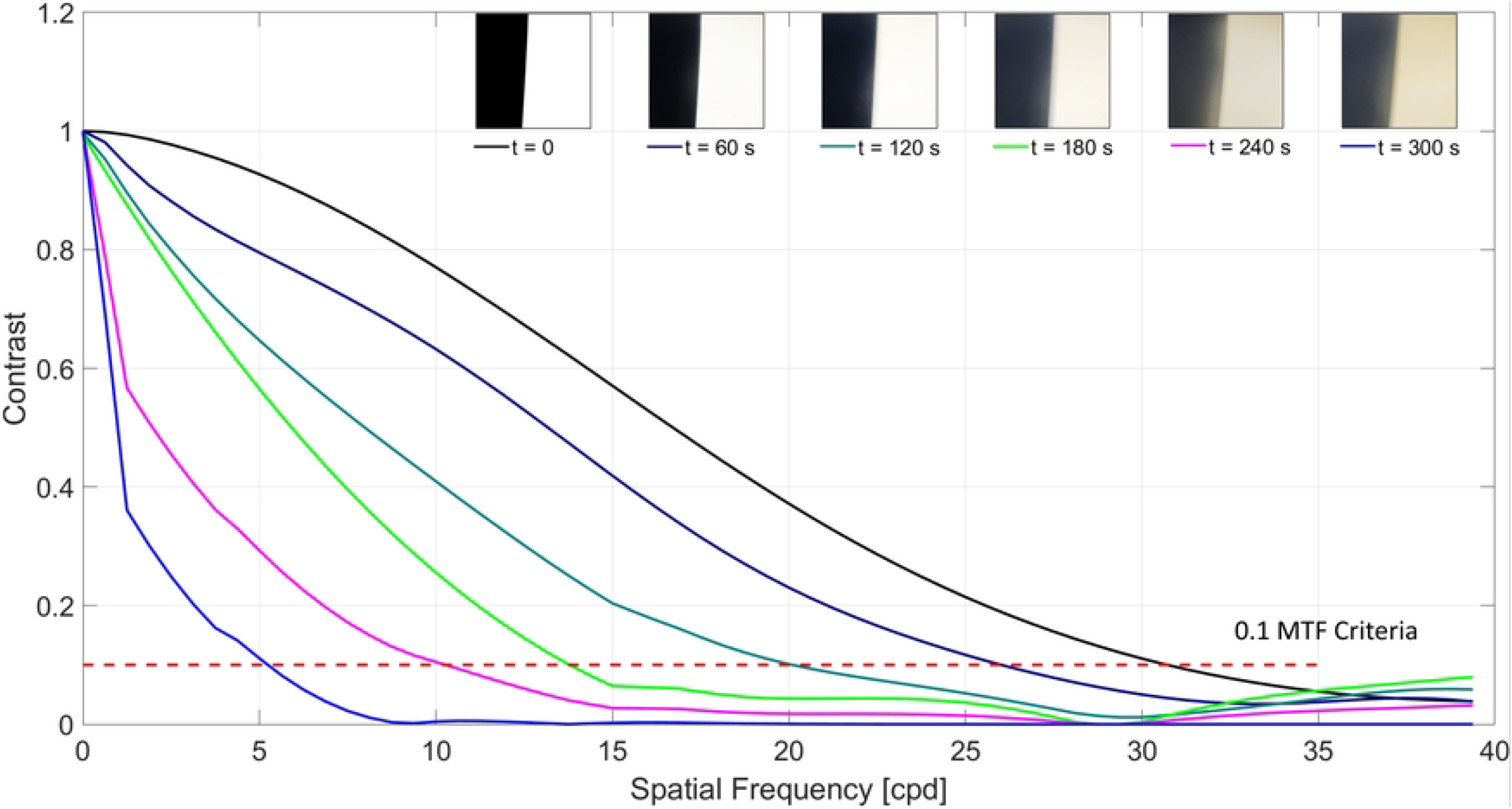
Nighttime capture through progressive opacification on IOL. The university courtyard is captured at night-time with the artificial eye model implanted with different opacification levels on the IOL at (A) 0 seconds, (B) 120 seconds, and (C) 240 seconds.

We also captured a sample reading chart from a close distance (35 cm) to see how cataracts would affect the reading function of a patient. The white arrow in the first capture of Fig 4 indicates 20/20 vision at a close distance. We can see that while without any cataracts the IOL performance barely reaches there, as the opacification starts and increases, the captures taken at 40 seconds intervals show that the visual acuity dramatically decreases. A more detailed analysis of the resolution of the process is done in the following parts of this study.

**Fig 4.**
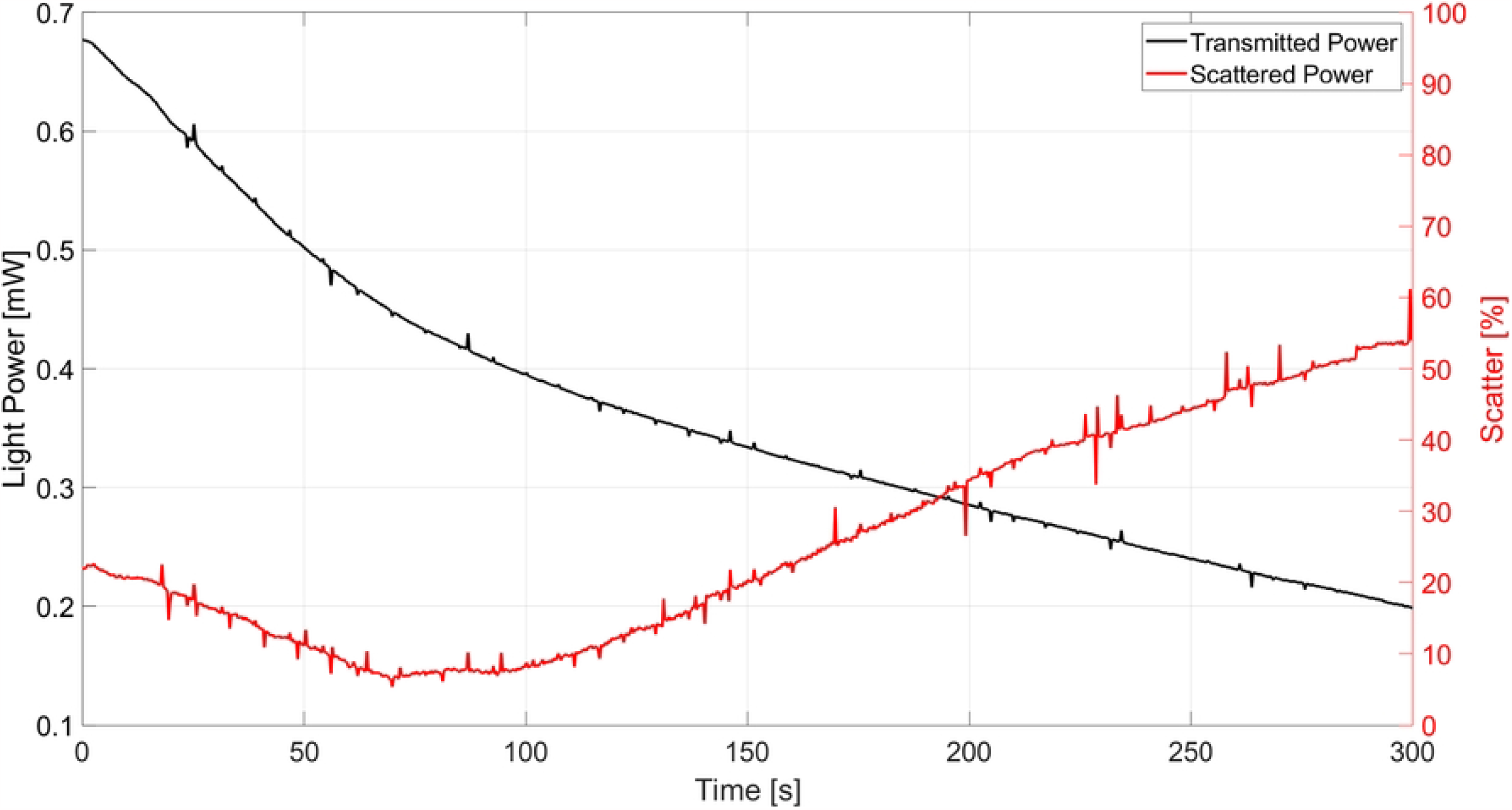
A sample close distance reading chart captured through an IOL with different opacification levels. The IOL is soaked in water for (A) 0, (B) 40, (C) 80, (D) 120, (E) 160, (F) 200, (G) 250, and (H) 300 seconds.

The holographic images were captured under different opacification levels as well as in a scenario where a pattern was made on the IOL, leaving a two mm-diameter area clear of any opaqueness (Fig 5). In the first scenario, the usual opacification steps were applied to the IOL. As the opacification proceeded, the holographic image was captured at equal intervals without changing the camera configurations (Figs 5A, 5B, and 5C). At first, the holographic image was clear, and the brightness of the image was sufficient. As the opacification proceeded, the quality and brightness of the holographic image decreased, and the blurriness of the letters increased, decreasing the resolution. In the second scenario, the holographic display sent the image through the non-opaque area of the IOL, and the holographic image was captured (Fig 5D). As it was observed, the capture quality and brightness were not as high as at the start,t where there was no opacification (Fig 5A), but the brightness and quality were relatively high compared to the increased opacification levels (Figs 5B and 5C).

**Fig 5.**
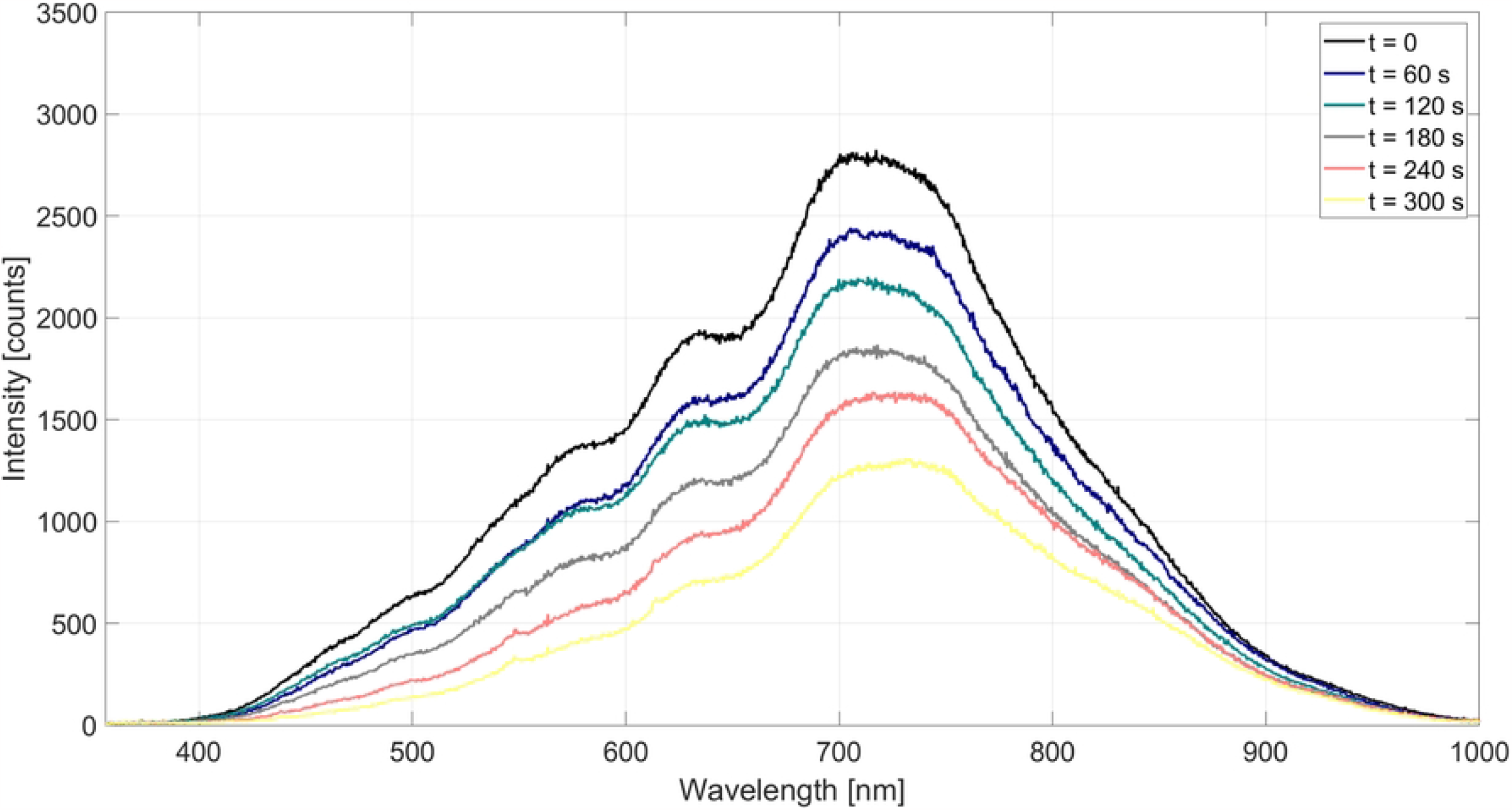
Holographic displays capture the Snellen reading chart. As the opacifications corresponding to soaking times (A) of 0 seconds, (B) of 150 seconds, and (C) of 300 seconds increased, the captures showed a decreasing brightness as well as a degradation of quality. (D) The capture of holographic display through a clear part of the IOL with cataract regions around it gave less quality compared to (A) but better quality compared to (B).

### Objectively Measured Parameters

Several characterizations were done to objectively understand the opacification process and how it degraded the vision through the IOL. The processes discussed in the methods section all yielded valuable results for us to understand if our surface opacification was correctly mimicking cataracts.

### MTF Charts

The MTF plots from the slanted edge captures were plotted as a function of spatial frequency for comparison (Fig 6). Thanks to the time dependence of the opacification, each capture represented different levels of opacification. At t = 0, there is no opacification on the IOL yet, and we can observe that its resolution reaches around 31 cpd, which is close to 20/20 visual acuity in a Snellen chart, when the MTF criteria is chosen as 0.1. As the opacification increases, with 60 seconds intervals, we can see that the achievable resolution decreases to 26, 20, 14, 10, and finally 5 cpd, respectively. Since 5 cpd is already a very low-quality resolution, the test wasn’t continued after 300 seconds. If the trend of resolution decrease is evaluated, we can say that there is nearly a linear decrease as a function of time; after 180 seconds, the quality of vision becomes very low.

**Fig 6.**
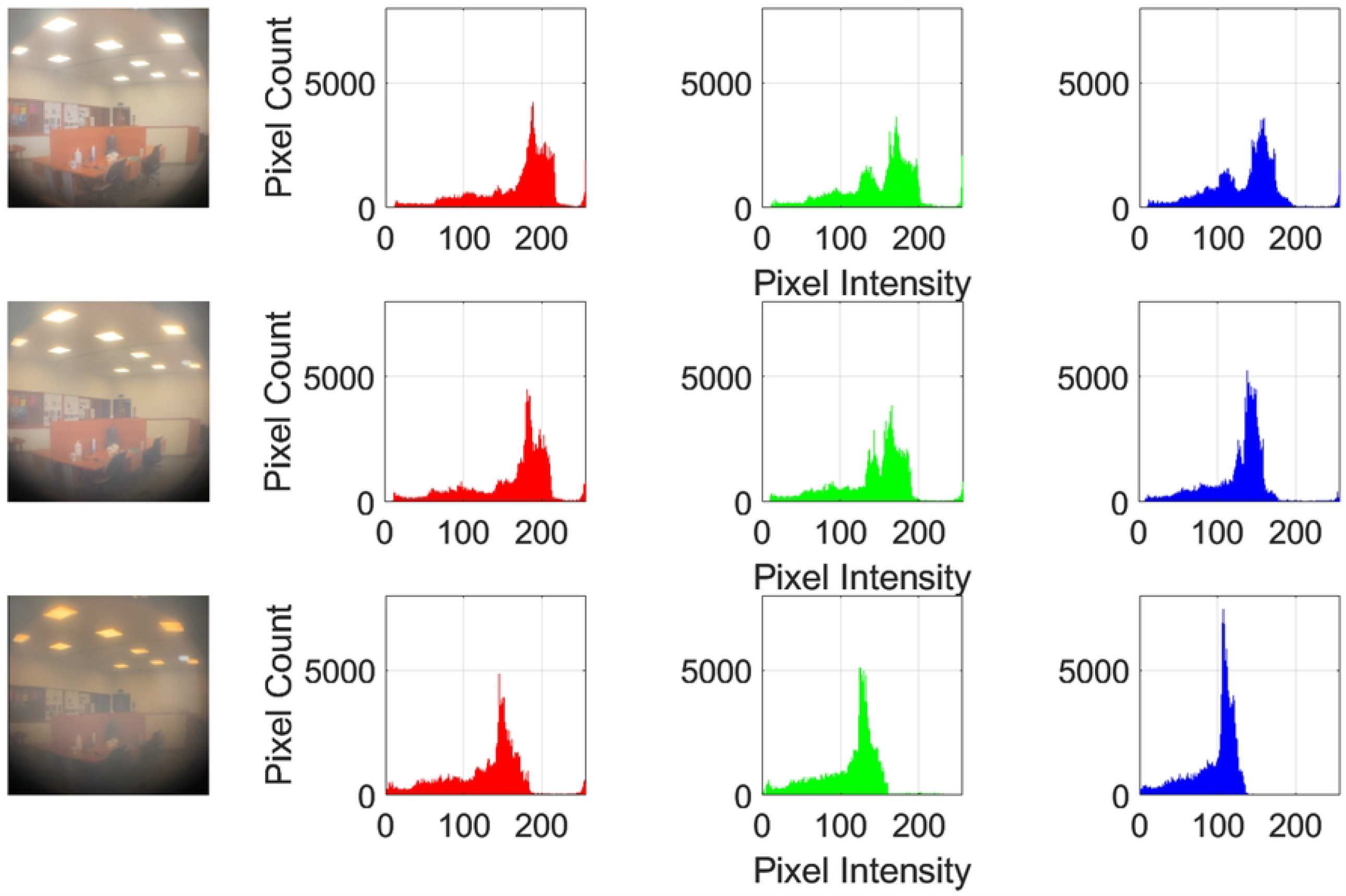
MTF charts obtained through the analysis of slanted edge captures through the IOL with different opacification levels from 0 to 300 seconds with 60 seconds intervals. As the opacification increases, it is seen that the achievable resolution decreases to 26, 20, 14, 10, and finally 5 cpd, respectively, with a 0.1 MTF criteria.

The transmitted light and the percentage of how much of it is scattered were plotted together as a function of time to see how they change with progressing opacification (Fig 7). As can be observed, the transmission starts at around 0.6 mW and decreases to 0.2 mW at the end of 300 seconds. Whereas scattering starts around 20 percent and goes up to 50% with passing time. This clearly shows that with increasing opacification on the IOL surface, the scattering of light increases as expected.

**Fig 7.**
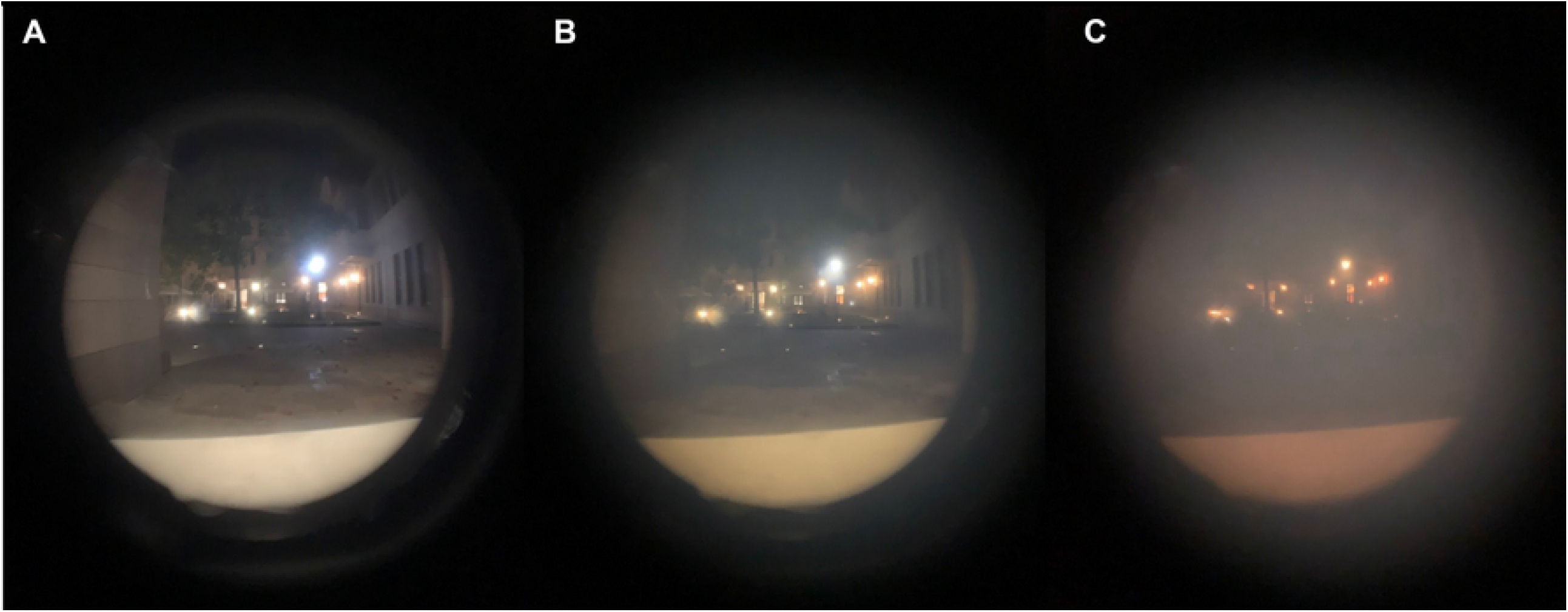
Transmission measurements of a 530 mW light source and the scattered percentage of transmitted light were plotted on the same graph as a function of time. As the IOL was soaked in the water cell, with passing time, the measured transmitted light decreased from 0.68 mW to 0.2 mW. The scattering of this transmitted light, in contrast, started around 20%, mildly decreased to 10%, and then sharply increased to 50%.

The results from the spectroscopy gave us valuable insights (Fig 8). It was seen that, at all wavelengths, a similar degradation could be observed. The important aspect of these results was that some wavelength intensities decreased more than others. For example, when the wavelengths around 700 nm are observed, we see that at the start the intensity count is around 2800, and it decreases down to 1200. This gives a 57% decrease in intensity. When we observe the intensity count around 600 or 800 nm wavelengths, we see that the starting point is around 1500. Then, as opacification increases, this value decreases to 500. This shows a decrease of around 33%, telling us that the peak around 700 nm is exposed to the highest intensity loss, and as we get away from that area, the percentage loss decreases. The results from spectroscopy measurements are further discussed in the S1 File.

**Fig 8.**
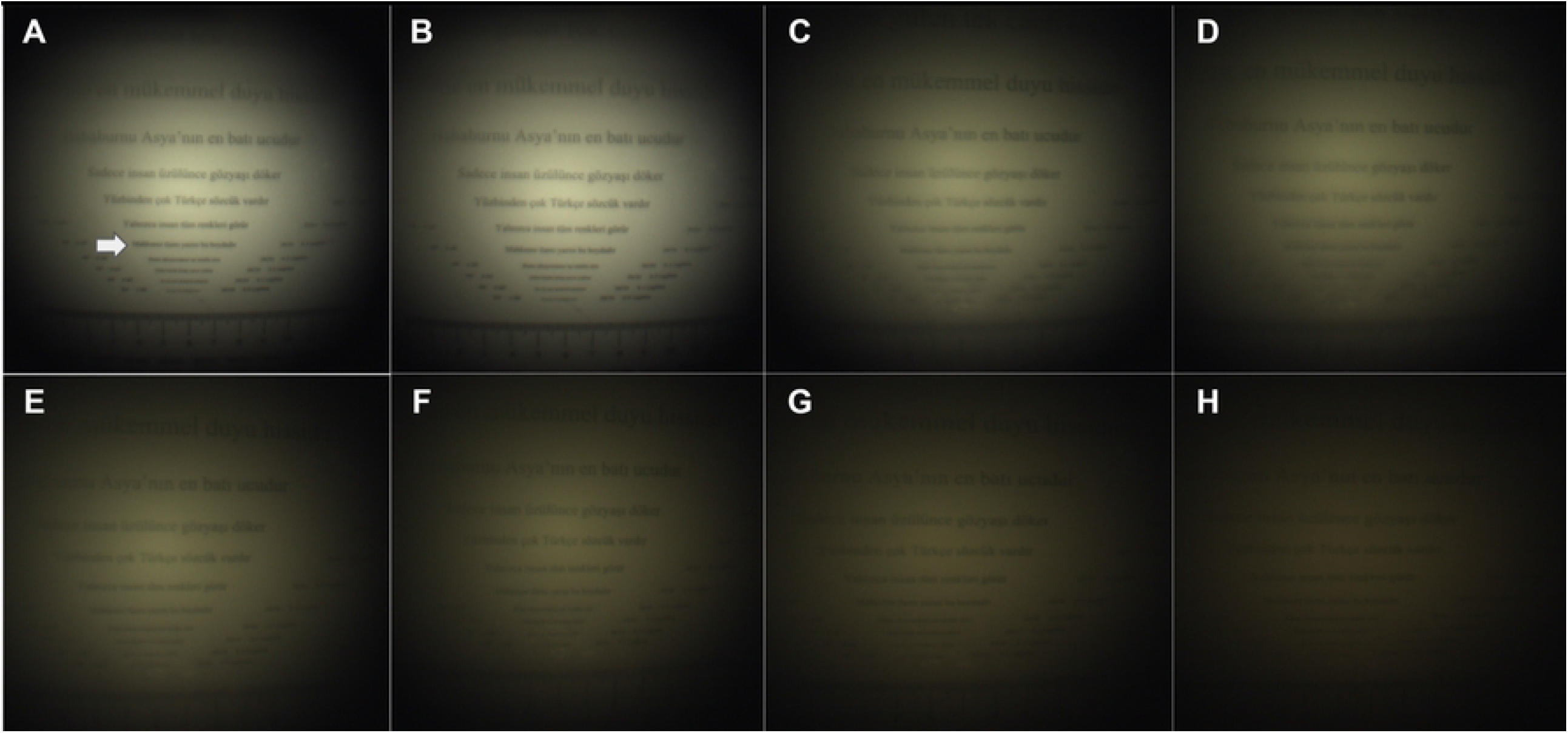
Spectroscopy measurements for different opacification levels corresponding to soaking time. As the soaking time increased, the intensity of all wavelengths decreased drastically, with the highest intensity loss occurring around 700 nm.

On the final part of the opacification process characterization, rather than a measurement, we conducted color histogram analysis on captures taken at different opacification levels (Fig 9). Three captures at 0, 120, and 240 seconds of opacification were analyzed to see how the color intensities were distributed. One general conclusion was that, as the opacification increased, the pixels shifted towards lower intensity values. A weighted average of each color’s percentage for each capture was also calculated. As seen, as the opacification increased, pixels tended to become more red and less blue, as expected. The percentage increase in green was negligible compared to the other colors. The pixels especially peaked around a pixel intensity of 100 at 240 seconds of soaking, losing nearly half of the initial intensity at blue and green and relatively less intensity in red.

**Fig 9.**
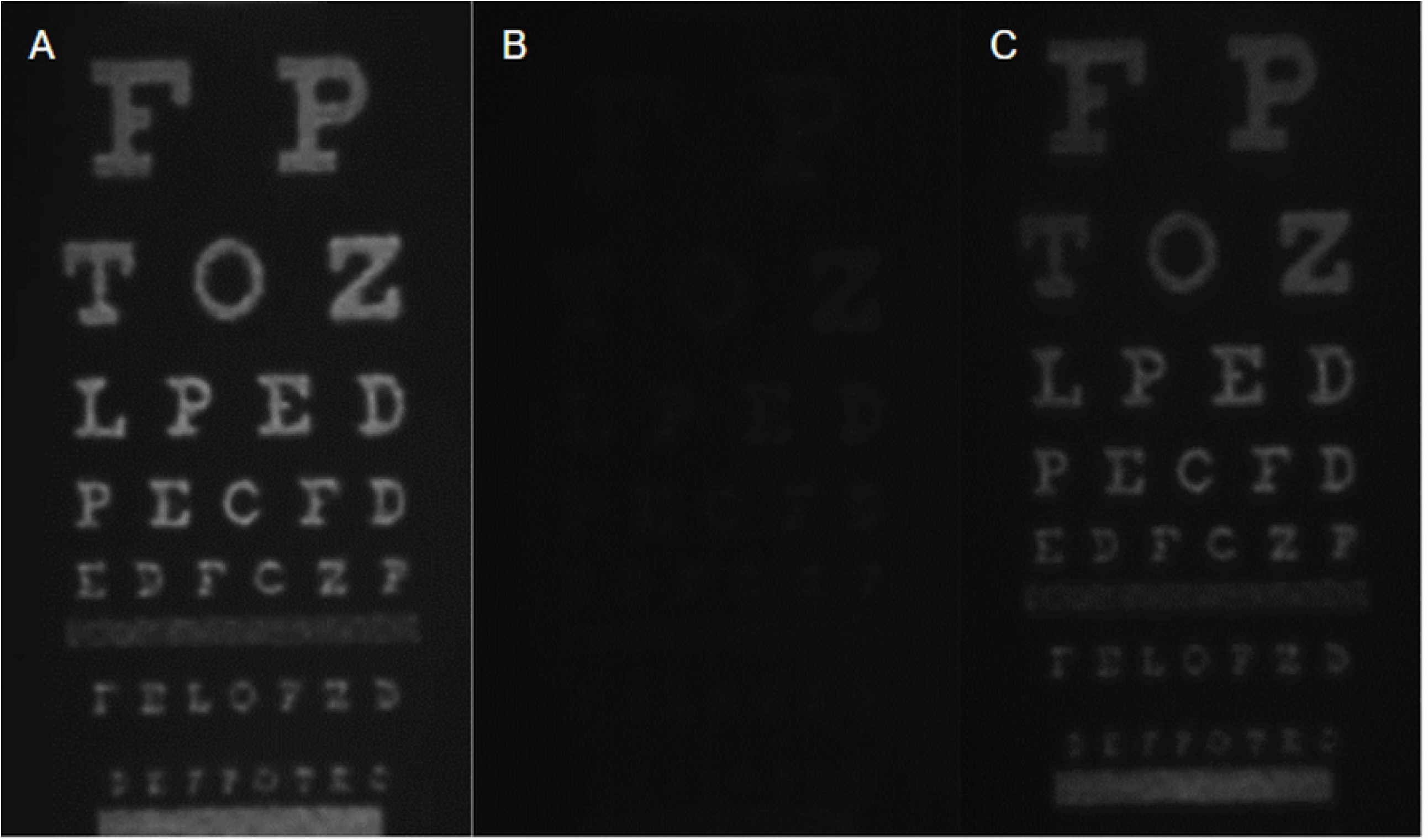
Color histogram and weighted average of pixels distribution to colors for opacification progress on IOL. At (A) 0 seconds, (B) 120 seconds, and (C) 240 seconds. From (A) to (C), as the opacification increases, the weighted average of red increases and blue decreases slightly. All colors show a tendency to shift to lower pixel intensities with the passage of time.

## Discussion

In this paper, we propose a simple method utilizing acetone and IOL to create a scattering surface (opaqueness on IOL) which mimics cataractous lens. Moreover, for the first time in literature, we combined this method with a powerful holographic display, in which we can pass any digital content through the opaqueness of the IOL.

Previous investigations replicated this opacification physically or digitally, using virtual reality, augmented reality, or optical setups. However, these virtual investigations were dependent on the subjective feedback of the cataract patients, and the single physical example was an irreversible process that would not be recreated on the same substance.

The advantage of this method over other methods is that it is a reversible process, so the opacification can be reversed at any time to repeat and observe the effects on vision. Furthermore, it clearly shows the effect of the level of opacification on vision. However, this process wears out the IOL as it is repeated, so the repetition limit is up to 20 times before the IOL starts to break apart. Another advantage of this method is that by partially applying the acetone to the opaque surface of the IOL, a cataract-like pattern can be created to simulate an even more realistic cataract case. Depending on how long the acetone is applied to the surface, while some areas have dense cataracts, others can have mild or no cataracts. Moreover, applying this method to selected surfaces of the IOL makes it possible to mimic different cataract or opacification cases and their progression.

All analyses showed the expected results. While the IOL without cataracts could show satisfactory visual performance by itself, as it was soaked in water after the acetone treatment and the opacification on its surfaces increased, there was a significant downgrade in its performance. The IOL choice was monofocal since the materials that were used in mono and multifocal lenses are the same, which does not perfectly represent the human crystalline lens in multi-depth scenarios. However, the primary concern of this study was to observe and understand the scattering and performance decrease overall, so it was not a significant issue.

Another unique point of this method was recreating the opacification process in a patterned sense. While there could be several options, such as masking the surface and treating only selected parts with acetone, cleaning the already opacified surface areas with alcohol was more practical. The impracticality of the masking lay in the small surface area, which would require very delicate work, causing our method to stray away from being simple.

We must mention one major limitation of our method. Since the chemicals are only applied to the surface and the opacification occurs on the surface, it does not physically represent natural cataracts, which occur inside the crystalline lens rather than just on its surface. This limitation did not affect the results much since the analyses showed that our method could successfully mimic cataract progression when the resulting visual qualities were observed.

An integral aspect of our study pertains to its potential clinical implications. With cataracts and PCO typically occurring slowly and patients often becoming aware of their condition only in severe cases, our approach holds promise as an educational tool within the realm of ophthalmology. This method could facilitate early diagnoses by demonstrating the progressive impact of opacification on vision, allowing for timely interventions that prevent a significant decline in patients’ quality of life.

## Conclusions

In this study, we tried to show the different levels of cataract on visual performance based on a novel acetone opacification technique that is combined with holographic display technology. We created a reversible and progressive opacification technique for IOLs using a unique chemical procedure utilizing distilled water and acetone. Using this technology, we recreated the slow decline of vision associated with various eye diseases. Unlike earlier irreversible approaches, our approach is reversible, allowing for repeated observations and analysis of the opacification process.

While existing studies have ventured into opacification simulation using augmented and virtual reality, our study introduces an experimental framework that bridges the gap between physical and virtual simulations. By replicating the progression of cataracts on IOL surfaces, we achieved a more realistic representation of opacification and preserved the ability to assess vision deterioration at varying stages. The versatility of this method further allowed us to recreate opacification patterns, simulating distinct cataract cases and their progression with controlled precision. Vision through areas with different opacification levels as well as clear parts surrounded by cataracts was assessed by using the programmable exit pupil patterns of our holographic display technology.

In conclusion, our study successfully introduces a reversible and adaptable method for simulating different levels of cataracts. The combination of our physical simulation approach, holographic displays, and objective analyses has produced compelling results that enhance our understanding of the visual implications of opacification. The insights garnered from this study will contribute to advancements in medical research and clinical practice, fostering greater awareness of opacification progression and early interventions for improved patient outcomes.

## Supporting information

**S1 Figure. Photodiode measurement setup**. Capture of the photodiode setup to measure transmission and scattering. An LED light source with a 530 nm wavelength was used. An aperture of 2 mm diameter then clips the incoming beam, and the collimating lens collimates it. The 2 mm diameter beam then goes through the wet cell and IOL with the cataractous surface, finally reaching the photodiode for measurement. For scattering, the difference in the setup was placing the photodiode a sufficient distance (6.66 cm) away from the wet cell. Hence, the IOL focused the beam on it, and an extra aperture just before the photodiode helped us eliminate the scattering light and measure only the focused light.

**S2 Figure. Spectrophotometer setup**. The spectroscopy measurements were conducted on this setup. A light source with a bandwidth of 100 nm to 1000 nm was used here. First, the light was passed through some ND filters, so the measurements were just below the saturated region. Then the light passed through the IOL with a cataractous surface and illuminated the spectrophotometer, which measured its intensity across wavelengths.

**S3 Figure. Holographic display setup**. The off-axis benchtop holographic setup has been meticulously designed, featuring a 520 nm central wavelength light-emitting diode (LED) as the light source. It incorporates essential components such as a spatial light modulator (SLM), an optical 4-f system, a magnifying lens, a tunable lens, an eyepiece lens, a wet cell (utilized for IOL positioning), and a camera. The SLM has a resolution of 1920 x 1080 pixels, and a 35-mm focal length lens is employed to collimate the LED light. Within the 4-f system, two convex lenses with 50-mm focal lengths are aligned alongside an iris diaphragm, strategically positioned in the Fourier plane. This diaphragm effectively acts as a spatial filter to eliminate unwanted diffraction orders originating from the SLM, allowing the desired holographic content to pass through. Incorporating a 50-mm focal-length lens, the holographic content is magnified for enhanced clarity. The tunable lens is essential for precisely adjusting the focus onto the hologram’s plane. To converge the content towards the IOL within the wet cell, the eyepiece lens is effectively employed. The context includes: a) a visual rendering of the optical setup for illustrative purposes, and b) details about the benchtop configuration utilized in our IOL experimentation.

**S4 Figure. IOL with patterned cataracts**. A small area on the cataractous surface on both sides of the IOL was cleaned with acetone to create a clear area. (A) Capture of the IOL before any procedure; (B) capture after the procedure of opacification and area clearing.

**S1 File. Normalized spectroscopy results and wavelength shift analysis**.

